# Regulatory divergence of flowering time genes in the allopolyploid *Brassica napus*

**DOI:** 10.1101/178137

**Authors:** D. Marc Jones, Rachel Wells, Nick Pullen, Martin Trick, Judith A. Irwin, Richard J. Morris

## Abstract

Polyploidy is a recurrent feature of eukaryotic evolution and has been linked to increases in complexity, adaptive radiation and speciation. Within angiosperms, such events occur repeatedly in many plant lineages. We investigated the role of duplicated genes in the regulation of flowering in *Brassica napus*. This relatively young allotetraploid represents a snapshot of evolution and artificial selection in progress. In line with the gene balance hypothesis, we find preferential retention of expressed flowering time genes relative to the whole genome. Furthermore, gene expression dynamics across development reveal diverged regulation of many flowering time gene copies. This finding supports the concept of responsive backup circuits being key for the retention of duplicated genes. A case study of *BnaTFL1* reveals differences in cis-regulatory elements downstream of these genes that could explain this divergence. Such differences in the regulatory dynamics of duplicated genes highlight the challenges for translating gene networks from model to more complex polyploid crop species.

---

Many economically important crops exhibit extensive gene multiplication as a result of recent or ancestral polyploidy^1^, for example wheat (*Triticum aestivum*)^2^, cotton (*Gossypium hirsutum*)^3^, and oilseed rape (OSR, *Brassica napus*)^4^. The presence of multiple copies of a gene relaxes natural and artificial selective pressures on any one individual copy, facilitating the emergence of novel gene functions^5^. The resulting increase in variation can be exploited to breed crop varieties with desirable phenotypes^6^. The presence of multiple orthologues, however, hinders efforts to translate knowledge of gene function and, in particular of regulatory networks, from model to crop species. This is a consequence of not knowing which orthologue, if any, retains the same function as the corresponding gene in the model species, whether ancestral functions have been partitioned between them, or if a novel function has been acquired^7^.

The evolutionary fate of gene copies arising from a gene duplication event has been studied in a range of species^8–11^. There are two main classes of gene duplication events: small scale duplications and whole genome duplications (WGD)^5,7,12–14^. These two types of duplication event can lead to different outcomes for gene copies^13^. Whilst gene redundancy has been reported to be evolutionarily unstable^7,15^, it is frequently observed^12,16–18^. A proposed driver for the retention of duplicate genes is the maintenance of gene dosage, known as the gene balance hypothesis^14,19–23^. Such dosage constraints may result if the gene product acts as part of a protein complex, where an incorrect stoichiometry of proteins can lead to the appearance of deleterious phenotypes^14^. WGDs maintain the original stoichiometry, resulting in duplicated, dosage sensitive gene orthologues being retained^14,20,23^. Conversely, small scale duplication of individual genes without their partners disrupts protein stoichiometry and disfavours gene retention^19^. Simulations of the dynamics of gene duplication events suggest that genes whose products form protein complexes, such as those associated with kinase activity, transcription, protein binding and modification, and signal transduction, are preferentially retained in the genome for longer when copied in whole genome relative to small scale duplications^19,24^. Data from a range of species are consistent with gene dosage balance^25–29^, including studies focusing on gene retention in the Arabidopsis genome^12,24^. In *Saccharomyces cerevisiae*, genes retained following a WGD are enriched for those that in diploids have haploinsufficiency or overexpression phenotypes, suggesting that the dosage of these genes is important^9^. One expectation of the gene balance hypothesis, illustrated in *S. cerevisiae*^20^, is that duplicated genes are more likely to be coregulated^20,23^. This co-regulation fits with the concept of buffering against stochastic effects in development^30,31^. Studying the regulation of duplicated genes can therefore provide clues for understanding their retention in the genome.

The *Brassica* genus contains several diploid crop species derived from ancestors that underwent a genome triplication event 5 to 28 million years ago^32–34^. OSR is an allopolyploid resulting from the interspecific hybridisation of two diploid species, *Brassica rapa* and *Brassica oleracea*^4^. An important agronomic trait for all Brassica crops is flowering time^35–38^, as different growing regions require varieties with very different phenologies. Flowering time has been extensively studied in the model species Arabidopsis^39–41^, revealing that flowering time genes are involved in multiple interactions and that many are transcription factors^41,42^. Thus, following the gene balance hypothesis, in a polyploid such as OSR, we would expect orthologues of Arabidopsis flowering time genes to have been preferentially retained relative to other genes in the genome, analogous to previous results that show preferential retention of genes involved with the circadian rhythm in paleopolyploid *B. rapa*^43^. That aspects of flowering time control are conserved between the Arabidopsis and OSR^37,44,45^ makes OSR an interesting and agronomically important model to investigate the evolution of gene function following gene multiplication.

Here we show that data from a transcriptomic time series (global gene expression in the first true leaf and shoot apex prior to and during the floral transition in OSR) support the prediction of preferential retention for flowering time genes in the genome (Figure 1). Through comparative gene expression and cluster analysis we demonstrate that the regulation of many flowering time gene homologues has diverged, suggesting this may be important for their retention. As an exemplar, using knowledge of cis-regulatory elements downstream of the Arabidopsis *TERMINAL FLOWER 1* (*AtTFL1*) gene, we identify sequence variation that correlates with regulatory differences observed for orthologues of *AtTFL1* in OSR. This case study highlights the importance of homologue expression dynamics in characterising gene regulation. The differences in *BnaTFL1* expression dynamics between homologues suggests that, in addition to proposed gene dosage effects, regulatory divergence may be important for gene retention.

**Figure 1.**
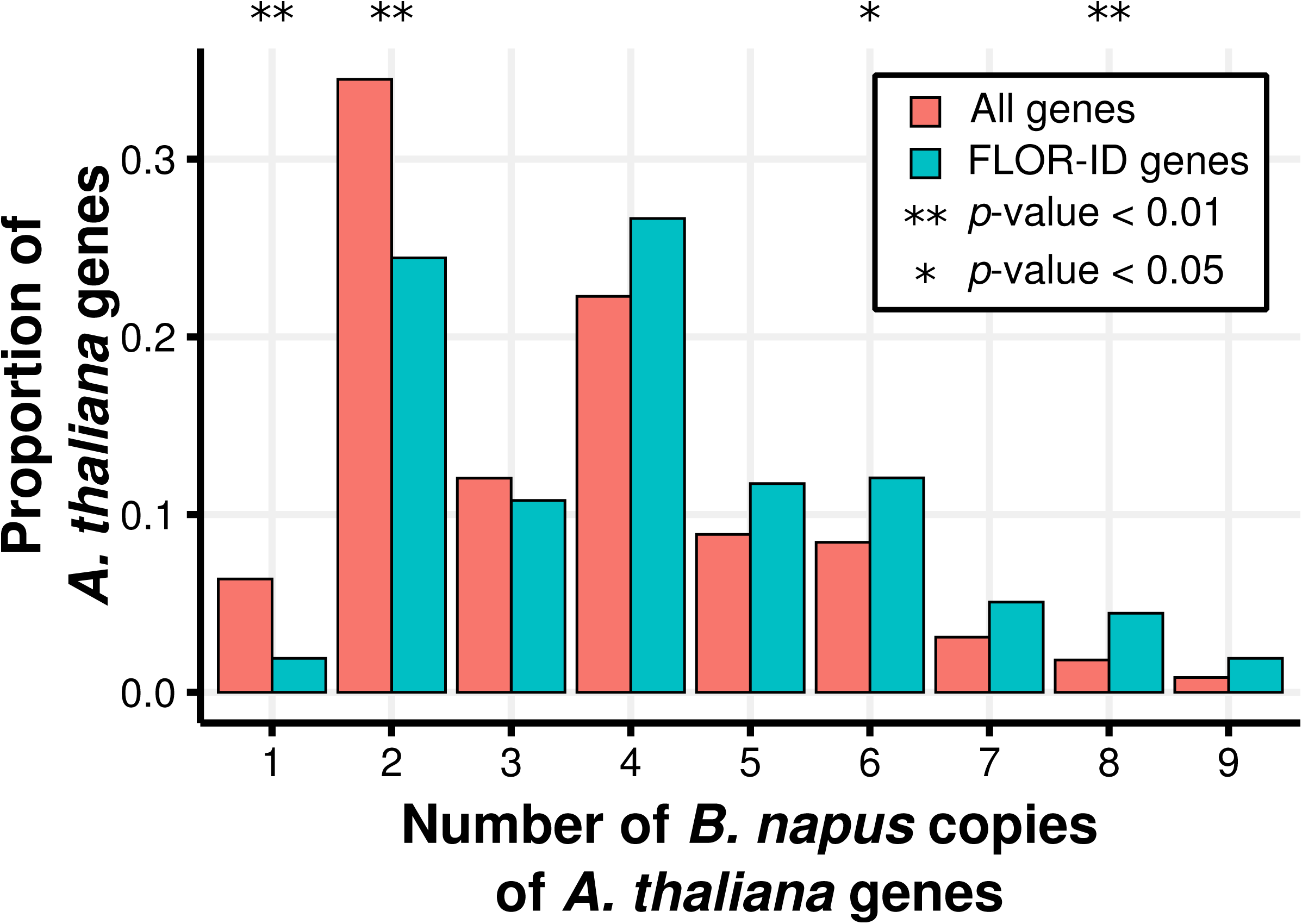
Arabidopsis flowering time genes have been maintained in the OSR genome at a higher copy number relative to other Arabidopsis genes. Annotated OSR genes were assigned to an Arabidopsis gene by taking the highest scoring BLAST result. The proportions were calculated by counting the number of Arabidopsis genes with a particular number of identified OSR copies and dividing by the total number of Arabidopsis genes represented by at least one gene in OSR. The FLOR-ID distribution is calculated using a subset of 315 Arabidopsis genes annotated as being involved with flower development or flowering time control in the FLOR-ID database^40^. False discovery rate corrected *p-*values were calculated by taking 1000 samples of 315 Arabidopsis genes from the 20882 represented in the All distribution. The mean and standard deviation of these samples were used to perform a two-tailed test of observing a proportion as extreme as the FLOR-ID value.

## Results

### OSR exhibits genome level expression bias across tissue types

Previous reports have demonstrated genome dominance in polyploids^46–48^. To test whether this is the case for OSR, we collected gene expression data through the vegetative to reproductive transition in a doubled haploid (DH) line derived from the spring OSR variety Westar (Figure 2). We compared global expression differences between the A and C genomes in the apex and the first true leaf across all time points (Figure 3; Supplementary Figure 1). We find that the A genome has a greater proportion of highly expressed genes than the C genome. Conversely, for genes showing very low expression we find the opposite relationship (Figure 3a). Similar distributions are found but are less pronounced when only OSR genes showing sequence conservation to annotated Arabidopsis genes are considered (Figure 3b) and when the sample is further restricted to OSR flowering time genes (Figure 3c). In contrast to the tissue-specific genome bias demonstrated in cotton^49^, our results are consistent across the two tissue types and throughout the time series (Supplementary Figure 1).

**Figure 2.**
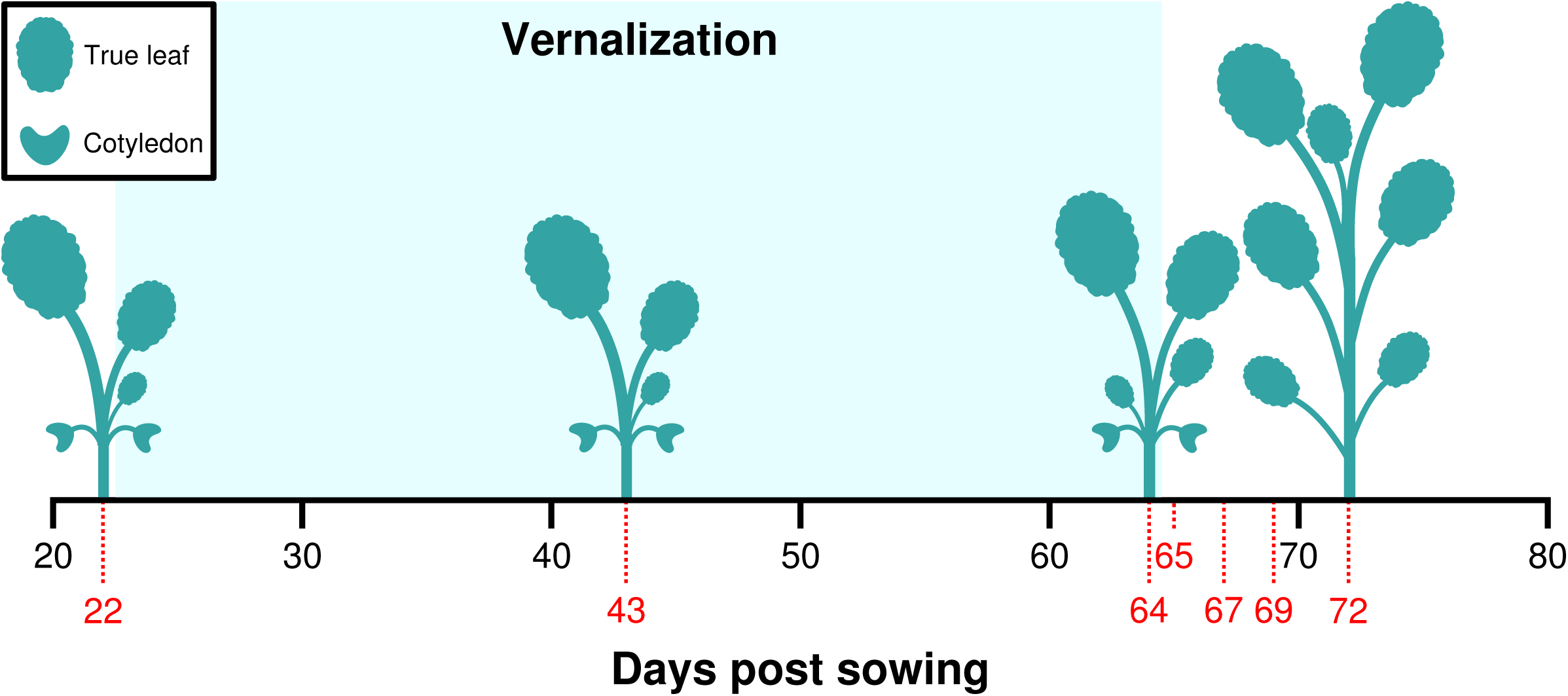
Tissue samples were collected for RNA-Seq at selected points through development. Plants were grown as detailed in the Methods. Tissue was sampled on the days indicated with red dotted lines and numbers. The plant silhouettes represent the approximate number of full leaves at the indicated points in development.

To investigate A and C genome expression at the gene level, we compared pairs of homoeologous genes that we identified using synteny and sequence similarity^4^. We classified a homoeologous pair as showing biased expression toward one genome if that gene has an expression level (measured in Fragments Per Kilobase of transcript per Million mapped reads, FPKM) at least two-fold higher than its homoeologue. At the individual gene level, biased expression was observed towards both genomes, but with 1.5 to 2.0 times as many genes showing bias towards the C rather than the A genome (16.9% towards the C genome relative to 9.7% towards the A genome in the apex, and 15.2% compared to 8.2% in the leaf; Table 1). This pattern is consistent with the findings of Chalhoub et al. (2014) and is maintained across all time points (Supplementary Table 2). The distributions of fold expression changes reveal that homoeologous gene pairs exhibiting a 2 to 8-fold change are primarily responsible for the observed bias (Supplementary Figure 2). Therefore, the homoeologue-level analyses reveal expression bias towards both the A and C genomes that are consistent across the tissue types tested and result in an absence of genome dominance (Supplementary Table 2). At the whole genome level, however, we observe a bias towards the A genome. This discrepancy may be due to genes with low expression levels tending to lack homoeologue pair information (Supplementary Figure 3). Alternatively, this bias may reflect a known higher incidence of homoeologous exchanges in which C genome copies of individual genes are replaced by their A genome counterparts^50^.

**Table 1.** Number of genes expressed 2-fold higher than their homoeologue for all homoeologue pairs. Homoeologue pairs^*4*^ were determined and filtered at each time point for those which both had expression levels above 2 FPKM. The number and percentage of these genes expressed 2-fold higher than their homoeologue is given. Despite some pronounced differences at the gene level, at the genome level the overall expression change is modest: The geometric mean of the fold difference of the C genome gene relative to the A genome homoeologue for all homoeologue pairs is 1.12 in the apex and 1.11 in the leaf.

| Days post<br>sowing | Apex |  |  | Leaf |  |  |
| --- | --- | --- | --- | --- | --- | --- |
|  | Both | A genome 2- | C genome 2- | Both | A genome 2- | C genome 2- |

**Table 1 – Number of genes expressed 2-fold higher than their homoeologue for all homoeologue pairs.**
|  | expressed | fold higher | fold higher | expressed | fold higher | fold higher |
| --- | --- | --- | --- | --- | --- | --- |
| 22 | 7313 | 596 (8.1%) | 1113 (15.2%) | 6294 | 620 (9.9%) | 1066 (16.9%) |
| 43 | 7389 | 597 (8.1%) | 1132 (15.3%) | 6176 | 626 (10.1%) | 1133 (18.3%) |
| 64 | 7325 | 602 (8.2%) | 1085 (14.8%) | 6307 | 597 (9.5%) | 1021 (16.2%) |
| 65 | 7243 | 609 (8.4%) | 1120 (15.5%) | 6182 | 601 (9.7%) | 993 (16.1%) |
| 67 | 7299 | 601 (8.2%) | 1135 (15.6%) | 6257 | 603 (9.6%) | 1046 (16.7%) |
| 69 | 7342 | 594 (8.1%) | 1130 (15.4%) | - | - | - |
| 72 | 7449 | 612 (8.2%) | 1119 (15.0%) | 6237 | 601 (9.6%) | 1054 (16.9%) |

**Figure 3.**
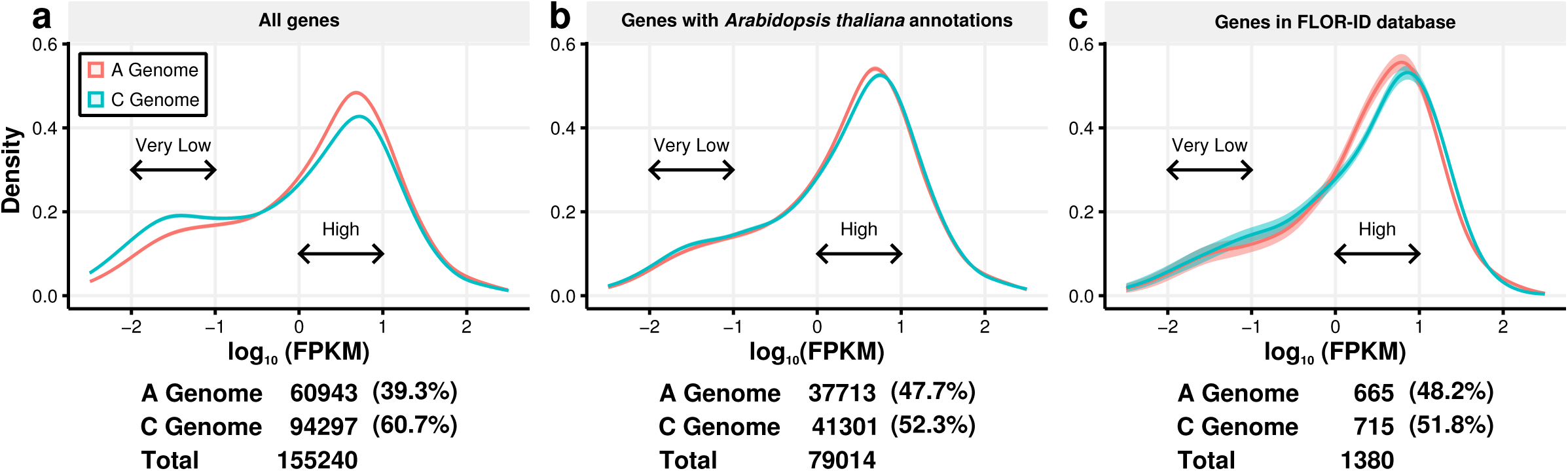
The A and C genomes of OSR show different patterns of gene expression. Density plots of transformed expression levels (log_10_(FPKM)) calculated using different gene subsets. The expression data was sampled 1000 times using a Gaussian error model. The density plot of log_10_(FPKM) values was calculated for each sample. The mean density and the 95% confidence interval estimated using the 1000 samples is displayed. Tabulated below each density plot are the number of OSR genes used to calculate the density plot, separated by their genome of origin. The data used to generate the density plots consisted of expression data from: **a** all annotated OSR genes, **b** OSR genes that show sequence conservation to an annotated Arabidopsis gene, and **c** OSR genes that show sequence conservation to an annotated Arabidopsis gene that is present in the FLOR-ID database^40^. These plots are generated using apex expression data from the time point taken at day 22, but are representative of the density plots obtained for all time points across both tissue types sampled (Supplementary Figure 1).

### OSR expresses a higher number of flowering time gene homologues relative to the whole genome

To test the prediction that flowering time genes are preferentially retained relative to the whole genome (Figure 1), we evaluated whether this was the case for genes expressed during the floral transition. A gene was considered to be expressed if the maximal expression level during the developmental time series was equal to or exceeded 2 FPKM, with leaf and shoot apex tested separately. We assessed the distributions of annotated (Figure 4a) and expressed OSR flowering time genes (Figure 4b and 4c). In both leaf and shoot apex (Figure 4b and 4c), a shift towards the expression of a higher number of flowering time gene copies relative to the whole genome can be observed. To test whether this observation was caused by the retention of circadian genes, as has been reported in *B. rapa*^43^, we repeated this analysis after removing this set of genes and found that the pattern remained (Supplementary Figure 4). This confirms the preferential retention of flowering time genes in OSR and suggests that the multiple orthologues of Arabidopsis flowering time genes retained in the genome could be functional.

**Figure 4.**
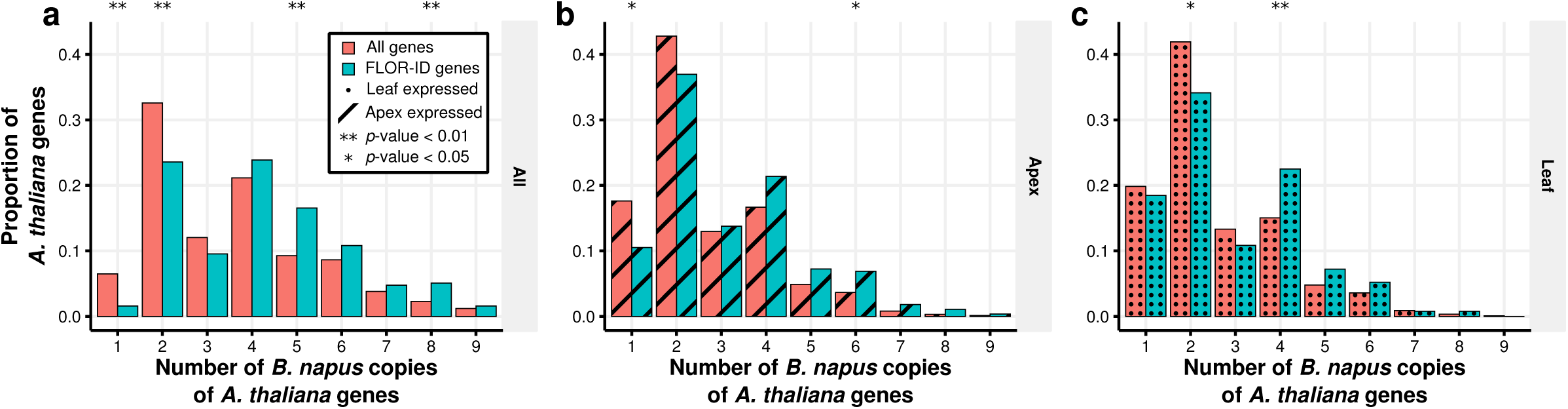
Multiple OSR flowering time gene homologues are expressed during the floral transition. The proportions of Arabidopsis genes that have particular numbers of homologues identified and expressed in OSR. OSR genes were considered to be expressed if their maximal expression level within a tissue across the time series was above 2.0 FPKM. False discovery corrected *p*-values are computed in the same way as Figure 1 using subsets of genes. **a** OSR genes that show sequence conservation to an annotated Arabidopsis gene. **b** OSR genes expressed in the apex tissue that show sequence conservation to an annotated Arabidopsis gene. **c** OSR genes expressed in the leaf tissue that show sequence conservation to an annotated Arabidopsis gene.

### Analyses of gene expression differences reveals regulatory divergence of retained flowering time genes in OSR

Having shown that genes involved in the control of flowering time are retained as multiple homologues in the OSR genome we next investigated their regulatory control. We first examined global gene tissue specificity and found that of the 45,048 genes expressed across the developmental time series, 16% show apex specific expression and 11% show leaf specific expression, with the rest (73%) exhibiting expression in both tissues (Supplementary Figure 6). Focussing on annotated orthologues of Arabidopsis flowering time genes, 61% have at least one orthologue in OSR that is not expressed in the apex, compared to 69% in the first true leaf (Figure 5).

**Figure 5.**
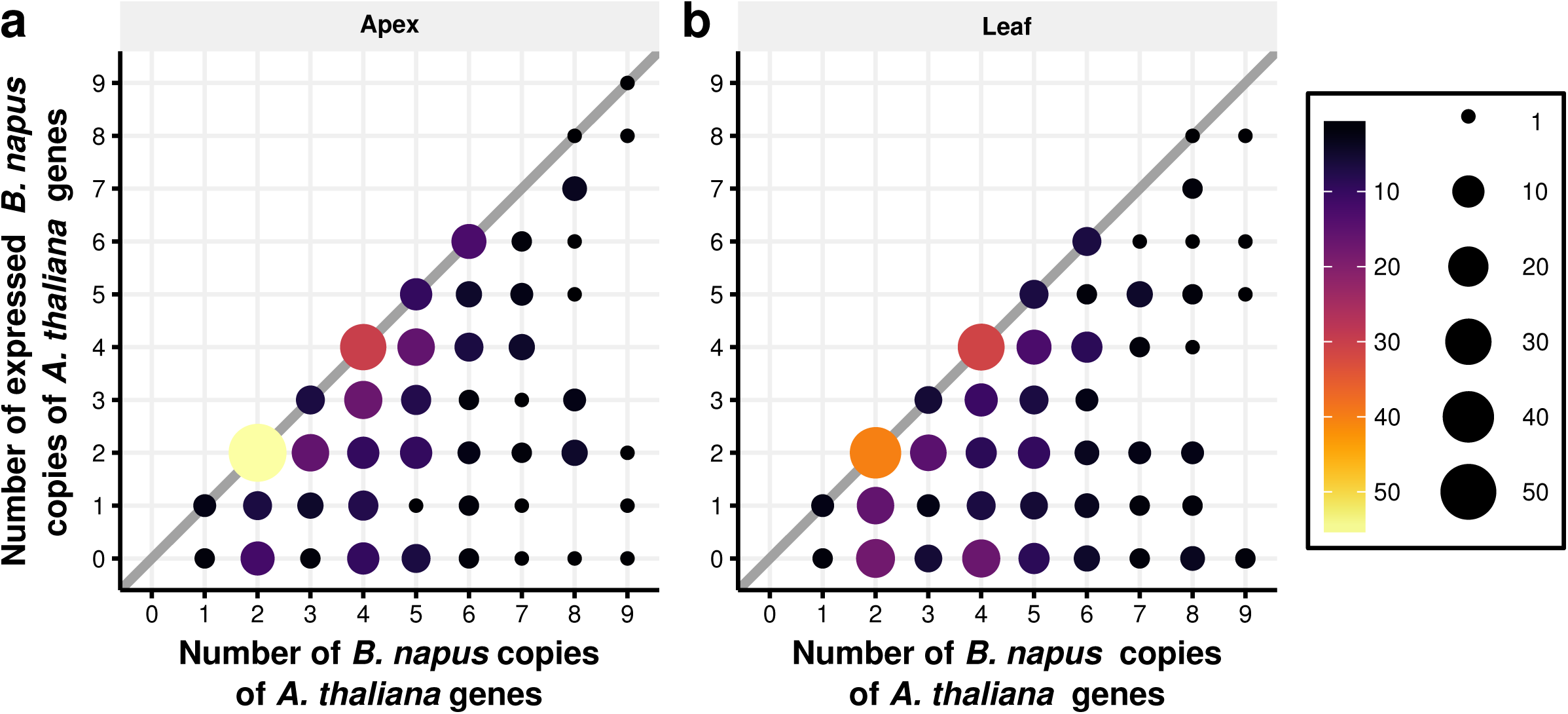
Not all annotated OSR orthologues of Arabidopsis genes are expressed. Expression data from the apex, **a**, and leaf, **b**, show that not all OSR copies of Arabidopsis genes were expressed in the developmental transcriptome time series. The size and colour of the circles indicate the number of data points at that position in the graph. The thick diagonal line indicates Arabidopsis genes that have OSR orthologues that are all expressed during the developmental transcriptome. Only OSR genes that show sequence conservation to an annotated Arabidopsis genes present in the FLOR-ID database^40^ were used to generate these results. A similar graph generated using all OSR genes that show sequence conservation to an annotated Arabidopsis gene is shown in Supplementary Figure 5.

We next used Weighted Gene Co-expression Network Analysis (WGCNA) to identify regulatory modules. WGCNA uses normalised expression data to cluster genes together based on their temporal expression profiles rather than expression levels *per se*. We used these cluster assignments to assess the regulatory control of flowering time gene homologues. Based on the premise of tight co-regulation of dosage-sensitive or functionally redundant genes^20,31^, our null hypothesis is that all OSR orthologues of an Arabidopsis flowering time gene will have similar expression patterns, leading to orthologues being in the same regulatory module (dashed lines in Figure 6). We found that most OSR flowering time genes (74% in apex, 64% in leaf) do not conform to this null hypothesis (Figure 6). Thus, analysis of both the overall level of expression in both leaf and shoot apex and WGCNA reveal regulatory divergence between retained homologues of flowering time genes in OSR, suggesting regulatory variation between homologues.

**Figure 6.**
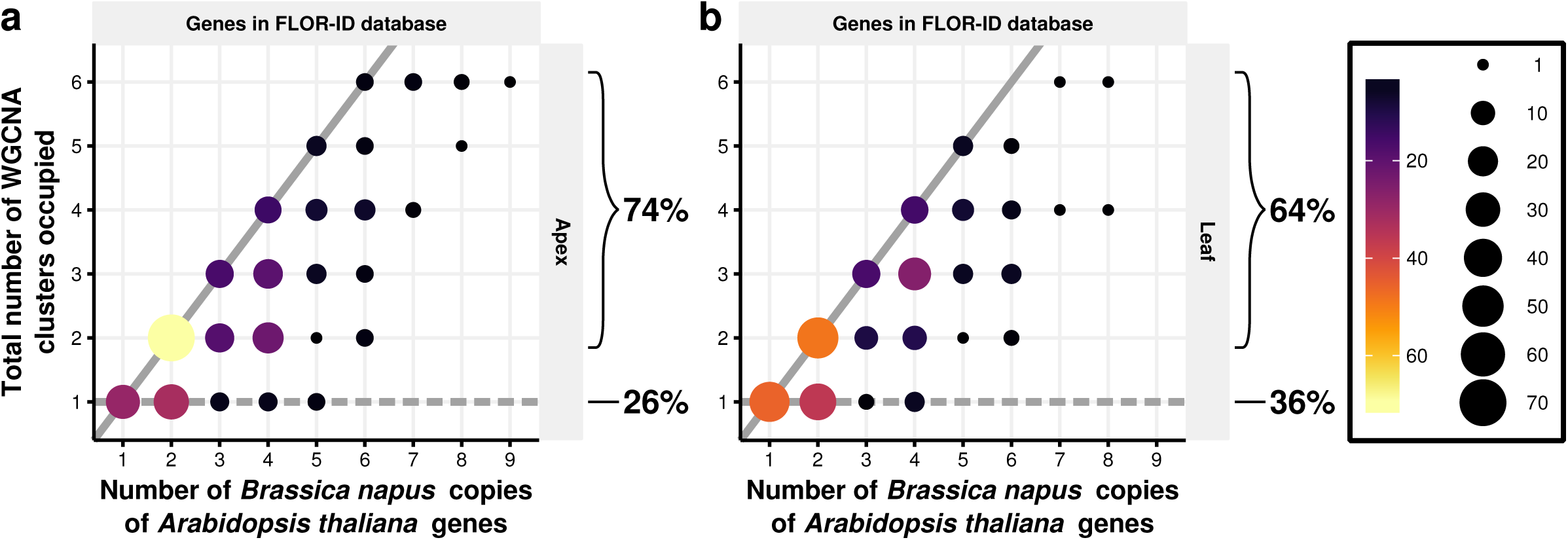
The majority of flowering time gene homologues in OSR are assigned to different regulatory modules. Regulatory module assignments for the apex, **a**, and leaf, **b**. The size and colour of the circles indicates the number of data points at that position in the graph. The thick lines on each graph represent two potential extremes. The dashed line represents the null hypothesis that all OSR copies of an Arabidopsis gene are assigned to the same WGCNA cluster. The solid line represents the Arabidopsis genes that have OSR copies that are each assigned to separate WGCNA clusters. The percentages indicated on the graph indicate the percentage of data points that agree, and the percentage that do not agree, with the null hypothesis. Only OSR genes with expression above 2.0 FPKM in at least one time point in the developmental time series and sequence conservation to an annotated Arabidopsis gene were used. A similar graph generated using all OSR genes that show sequence conservation to an annotated Arabidopsi*s* gene is shown in Supplementary Figure 7.

### Self-organising map based clustering captures different patterns of regulatory divergence for OSR orthologues of the flowering time genes *AtTFL1*, *AtFT*, and *AtLFY*

To further assess differences in regulation between gene homologues we analysed the divergence of expression over time. Whilst WGCNA assigns expression profiles to regulatory modules, the similarity between profiles is not quantified and genes that could be assigned to multiple regulatory modules are only assigned to a single module. Furthermore, WGCNA does not account for uncertainty in the RNA-Seq data in the assignment of regulatory modules. To address these issues, we employed a self-organising map (SOM) based sampling approach to assess expression profile divergence (Supplementary Figure 8). Figure 7a illustrates the five possible patterns of regulatory module assignment: (1) a *distinct* pattern of multiple regulatory modules with genes assigned to a single module; (2) a *gradated* pattern of multiple modules where gene membership of individual modules overlap; (3) a *unique* pattern (a special case of the *distinct* pattern) where each copy of a gene is assigned to a different module; (4) a *redundant* pattern where all genes are assigned to the same regulatory module; (5) a *mixed* pattern with some modules showing overlap in gene membership and others not. This approach allows us to robustly analyse expression similarity. Of 85 pairs of homoeologues expressed in the apex, 67 (79%) are found in the same regulatory module. In the leaf, 53 of 69 (77%) of expressed homoeologous pairs are found in the same module, with 29 of the co-regulated pairs being common between the two tissues (Additional File 1). The percentage of Arabidopsis genes with at least two expressed homologues in the apex (leaf) exhibiting each of the regulatory module assignments are 25% (26%) *distinct*, 9% (6%) *gradated*, 23% (23%) *unique*, 39% (33%) *redundant*, and 3% (6%) *mixed* (Supplementary Figure 8).

**Figure 7.**
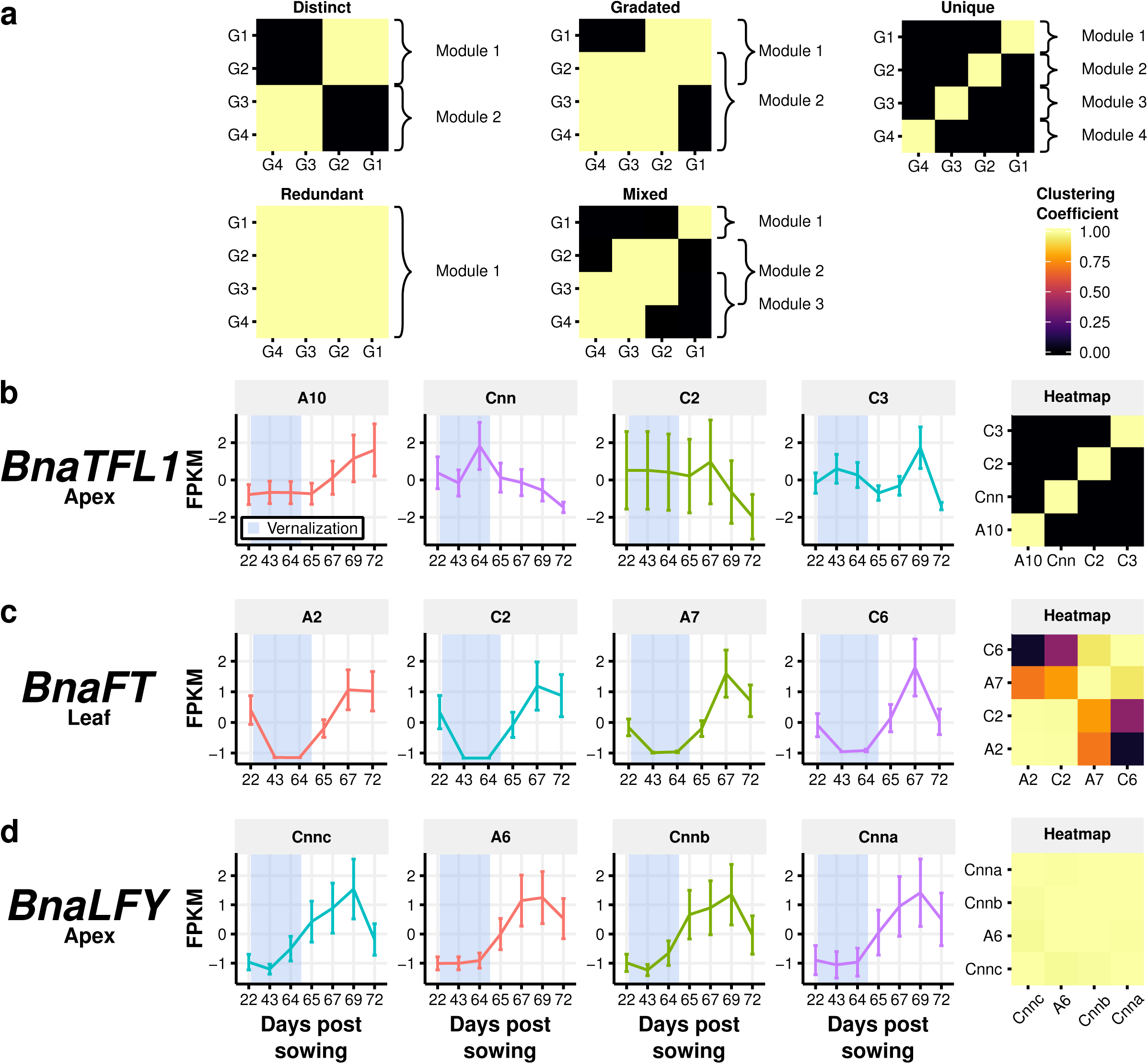
The OSR orthologues of *AtTFL1*, *AtFT*, and *AtLFY* show different patterns of regulation. **a** Representations of the five patterns of regulatory module assignment detected by the SOM based method. High clustering coefficients between two different genes indicates that those genes have similar expression traces. Clustering coefficients between a gene and itself represent how robustly a gene maps to the SOM. A *distinct* pattern indicates multiple regulatory modules being identified, with no gene occupying more than one module. A *gradated* pattern represents multiple regulatory modules being detected, but genes occupy multiple modules. *Redundant* patterns occur when only one regulatory module is detected, and all copies of a gene are assigned to that module. *Unique* patterns are a special case of *distinct* pattern where each copy of a gene is assigned to a different regulatory module. *Mixed* patterns consist of a mixture of *distinct* and *gradated* patterns, where the gene assignment of some modules overlap while others do not show overlap. When assessing the regulatory module assignment, gene copies that do not robustly map to the SOM are removed. **b, c** and **d** Expression traces across the developmental time series were normalised to a mean value of 0.0 FPKM and unit variance across the time series. The shading indicates time points during which the plants were grown in cold conditions. Regulatory module assignment heatmaps calculated using the SOM based method for the OSR copies of *TFL1*, *FT*, and *LFY* are also displayed. Both the expression traces and the clustering coefficients are apex derived for *TFL1* (**b**) and *LFY* (**d**) and leaf derived for *FT* (**c**).

To investigate further we chose three central Arabidopsis flowering time genes *AtLFY*, *AtFT* and *AtTFL1*. These genes form key hubs in the regulatory network responsible for the switch to flowering in rapid cycling Arabidopsis^51^. Each of these genes has four expressed orthologues in OSR with *BnaTFL1* and *BnaLFY* expressed in the apex and *BnaFT* expressed in leaf tissue. SOM analysis revealed that orthologues of *AtLFY*, *AtFT* and *AtTFL1* in OSR exhibit three different patterns of regulatory module assignment; *redundant*, *gradated* and *unique* respectively.

Homologues of *BnaLFY* exhibit a *redundant* pattern of regulatory module assignment, with each of the expression profiles in the apex showing low expression initially and an increase after the vernalisation period (Figure 7d), analogous to observations of *AtLFY* expression in Arabidopsis^52^. Co-regulation of *BnaLFY* homologues is consistent with the gene balance hypothesis^20,23^ and is supported by *AtLFY* displaying dosage sensitivity^52,53^.

The four *BnaFT* homologues exhibit a *gradated* pattern with two modes of regulation (Figure 7c). The expression of all homologues of *BnaFT* decreases during vernalisation and returns to pre-vernalisation levels when the plants are returned to growth in warm, long day conditions. The *BnaFT* expression profiles diverge at the final time point (day 72) with the A7 and C6 homoeologues showing a pronounced decrease in expression between days 67 and 72. The decrease in expression of *BnaFT.A7* is not as marked as that of its homoeologue, resulting in its assignment to both regulatory modules. The *BnaFT* homologues expressed in the leaf therefore exhibit a gradient of regulatory responses, with *BnaFT.A2* and *BnaFT.C2* having divergent expression traces relative to *BnaFT.C6*, but with *BnaFT.A7* showing similarities to all homologues.

OSR orthologues of *AtTFL1* are an example of *unique* regulatory module assignment with each of the four *BnaTFL1* genes assigned to different modules (Figure 7b). *BnaTFL1.A10* is expressed before and during cold with an immediate increase in expression when the plants are returned to growth in warm, long day conditions. *BnaTFL1.C2* also shows stable expression before and during cold but in contrast to *BnaTFL1.A10* decreases in expression when the plants are returned to warm, long day conditions. *BnaTFL1.C3* exhibits reduced expression levels post-cold with a transient peak of expression at day 69. The fourth homologue (mapped to the Darmor-*bzh* C genome and with greatest sequence identity to *BolTFL1.C9* from the EnsemblPlants database^54^) shows increased expression during cold followed by a steady decrease when plants are returned to warm, long day conditions. These four expression profiles are *unique* as shown in the clustering coefficient heatmap (Figure 7b). Homologues *BnaTFL1.A10* and *BnaTFL1.C3* exhibit expression profiles with the greatest similarity to *AtTFL1*^55^ as both show increasing expression during the floral transition.

*AtLFY*, *AtFT* and *AtTFL1* integrate environmental signals to determine the timing of the floral transition^56–60^. That individual orthologues of these genes in OSR show different patterns of regulatory module assignment suggests that the selective pressures acting on them are different, even though they belong to the same regulatory pathway in Arabidopsis. This result mirrors findings in Arabidopsis where it was found that less than half of gene pairs derived from the most recent duplication still retained significantly correlated expression profiles^12,26^.

### Patterns of intergenic sequence conservation surrounding *BnaTFL1* genes provide a potential explanation for the observed regulatory divergence

Downstream regulatory sequences of *AtTFL1* in Arabidopsis have been shown to be important for spatiotemporal control of expression^61^. We therefore investigated whether similar variation could explain the *distinct* pattern of regulation displayed by the four *BnaTFL1* orthologues. We analysed sequence conservation between OSR and Arabidopsis in the 5’ and 3’ intergenic regions surrounding *BnaTFL1*, identifying several conserved regions (Figure 8). Focussing on areas previously identified as *AtTFL1* cis-regulatory elements in Arabidopsis^61^, we find variation in the degree of sequence conservation between *BnaTFL1* orthologues (Figure 8a). Sequence conservation within regions II and IV of *BnaTFL1.A10* and *BnaTFL1.C3* suggests Arabidopsis*-*like cis-regulatory elements are present downstream of these genes. These *BnaTFL1* orthologues, that increase in expression during the floral transition, show high sequence conservation in region II. Conversely, *BnaTFL1.Cnn* and *BnaTFL1.C2,* which are not upregulated during the floral transition, lack sequence conservation in this region. Region II was found to be necessary for the upregulation of *AtTFL1* during the floral transition in Arabidopsis^61^, which correlates with this result. Region IV may also be involved in the observed expression trace divergence between *BnaTFL1* homologues, as this region was found to be important for the expression of *AtTFL1* in the inflorescence meristem.

**Figure 8.**
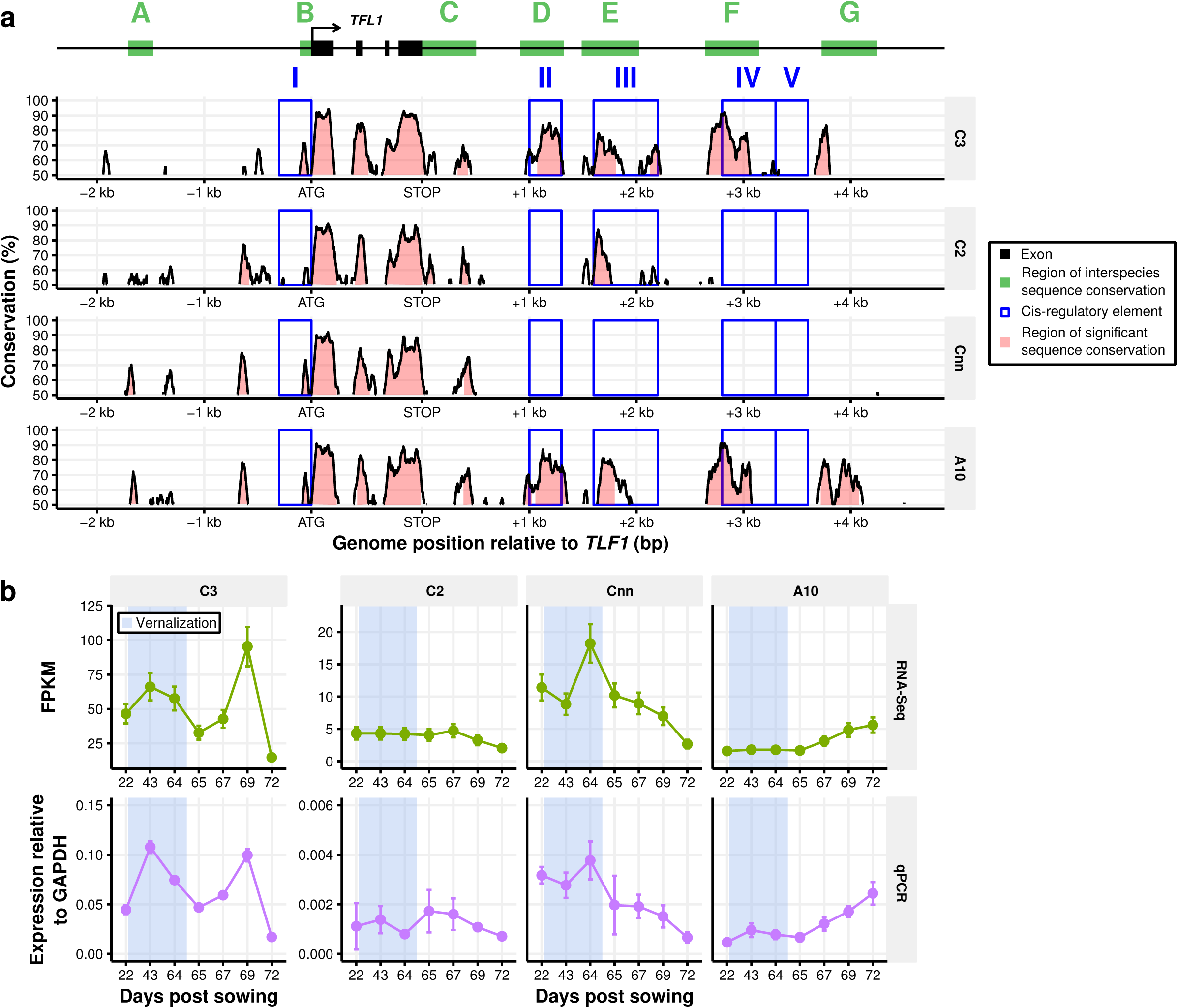
Sequence analysis reveals that cis-regulatory modules identified in Arabidopsis are not present downstream of some copies of *TFL1* in OSR. **a** The degree of sequence conservation between the OSR copies of *TFL1* and *AtTFL1*. Sequence alignment and conservation calculations were performed using the mVISTA server^88,89^ with a sliding window size of 100bp. The seven regions of high interspecies sequence conservation (green bars) and the five cis-regulatory regions (blue boxes) identified^61^ by Serrano-Mislata et al. are shown relative to the *AtTFL1* gene model (black bars). The labelling of these regions follows the same conventions as the previous study. The pink shaded areas under the sequence conservation curves are regions above 70% sequence conservation. Genomic position upstream and downstream of the *TFL1* gene copies are given relative to the ATG and STOP codon sites respectively. **b** The unnormalised expression traces for the *BnaTFL1* genes determined through RNA-Seq and RT-qPCR. The expression values calculated for RT-qPCR are normalised to *GAPDH* with the error determined from two biological replicates (Methods).

Sequence conservation within region III is below 50% in *BnaTFL1.Cnn*, whilst for the other three homologues it is 81%, 87%, and 78% for *BnaTFL1.A10*, *BnaTFL1.C2*, and *BnaTFL1.C3,* respectively. Interestingly, the range of significant sequence conservation in *BnaTFL1.C2* (154 bases) and *BnaTFL1.A10* (162 bases) is decreased compared to that of *BnaTFL1.C3* (273 bases), potentially suggesting the cis-regulatory elements in the former two copies are incomplete.

Serrano-Mislata et al. (2016)^61^ identified additional regions conserved across species that were not experimentally implicated in the regulatory control of *AtTFL1* (green shading in Figure 8). We observe sequence divergence in one of these regions, region G. Interestingly it is *BnaTFL.A10* and *BnaTFL1.C3,* which exhibit expression profiles most like that of *AtTFL1,* that show sequence conservation in this region. *BnaTFL1.A10* exhibits high sequence conservation relative to Arabidopsis across this entire region, while *BnaTFL1.C3* shows conservation over ∼50% of the region. As with regions II and IV, *BnaTFL1.C2* and *BnaTFL1.Cnn* lack conserved sequence in region G. We also identified a region of conservation not annotated in the previous analysis of *AtTFL1* cis-regulatory elements. This region, situated ∼600 bp upstream of the transcription start site of *AtTFL1,* shows ∼80% sequence conservation relative to Arabidopsis in *BnaTFL1.A10*, *BnaTFL1.C2* and *BnaTFL1.Cnn*. In *BnaTFL1.C3*, sequence conservation in this region is ∼55%.

To confirm the expression differences we observe between the *BnaTFL1* orthologues we performed copy-specific RT-qPCR across the developmental time series (Figure 8b). The RT-qPCR results show good correspondence with the RNA-Seq results, confirming our findings. Thus, using sequence conservation we determine the presence/absence of cis-regulatory elements downstream of the *BnaTFL1* genes that may confer similar regulatory control in OSR as in Arabidopsis. *BnaTFL1* orthologues contain different combinations of cis-regulatory elements, which have the potential to underlie the divergent expression traces they exhibit.

## Discussion

WGD events are thought to have occurred in most, if not all, angiosperm lineages^62^ and are well documented in the Brassicaceae^33,63^ Whole genome triplication^32–34^ and interspecific hybridisation events^4^ have resulted in extensive gene multiplication in Brassica species relative to the Arabidopsis lineage. WGD is considered a driving force in angiosperm diversification^64^, introducing genetic redundancy and allowing the evolution of novel gene function and new interactions, leading to neo- and subfunctionalisation. WGDs are usually followed by a process of “diploidisation”^65^ that includes genome downsizing^66^, chromosome rearrangement and number reduction^67^, and gene loss^68^. So, whilst many additional gene copies gained from WGD are likely to be lost over time, the analysis of genomic sequences has revealed that a significant number of duplicated genes are nevertheless present in the genomes of many species^12,16–18^. For instance, in the Arabidopsis lineage around 30% to 37% of homoeologous gene duplicates have been retained^25,69^. Based on such observations, modelling studies have determined conditions under which duplicated genes can become evolutionary stable^14,30^. These ideas have given rise to the gene balance hypothesis, which states that dosage sensitive genes are preferentially retained in the genome after WGD, but tend to be lost after local duplication events^20,23^. Kinases, transcription factors and proteins that form part of a complex fall into this category. From the gene balance hypothesis, we might therefore expect that highly networked genes such as those that regulate flowering time^40,41,70^ have been preferentially retained in the genome.

This study determines the expression profiles of OSR genes prior to and during the floral transition. We compared expression profiles across development to infer whether orthologues of Arabidopsis flowering time genes retain similar patterns of regulation. Whilst our analysis reveals that a significant proportion of duplicated genes in OSR have divergent regulation (Figures 5 and 6, Supplementary Figures 8b and 8c), it shows that the more recently combined homoeologues are frequently found in the same regulatory module (79% in the apex and 77% in the leaf). The finding of homoeologues tending to be co-regulated in allotetraploid OSR is intriguing, given the comparatively recent origin. An analysis of 2,000 pairs of paralogous genes in *Gossypium raimondii*, resulting from a 5-to 6-fold ploidy increase ∼60 Mya, revealed more than 92% of gene pairs exhibited expression divergence^71^. Most of these gene pairs show complementary expression patterns in different tissues, consistent with the idea of responsive backup circuits^15,31^. It is therefore tempting to speculate that regulation of homoeologues in OSR is still in flux with near-complete divergence a likely consequence of “diploidisation” across much longer timeframes. This hypothesis is supported by the finding that in recently synthesised allotetraploid cotton, most homoeologues display similar expression patterns across multiple tissue types^49^ while in allotetraploid upland cotton (*G. hirsutem;* which arose 1-2 Mya) 24% of homoeologues show diverged expression patterns. Recent genomic studies also support the idea that the OSR genome is in flux^50,72^, potentially in response to artificial selection for agronomically important traits.

Gene expression can be controlled through a range of mechanisms. This study highlights the potential role cis-regulatory elements may play in the divergence of gene regulation. Expression divergence of *AtTFL1* orthologues in OSR correlates with the presence and absence of sequence conservation within regions downstream of the gene. Serrano-Mislata et al. (2016) identified these regions as cis-regulatory elements and dissected their roles in the spatiotemporal regulation of *AtTFL1*^61^. *AtTFL1* expression dynamics exhibited by Arabidopsis mutants lacking the identified cis-regulatory elements show striking similarities to those of *BnaTFL1* orthologues lacking sequence similarity to the elements. This suggests conserved function of cis-regulatory elements between Arabidopsis and OSR and highlights that such variation can potentially drive the regulatory divergence of gene homologues. Although the patterns of sequence conservation downstream of *AtTFL1*^61^ are retained in OSR orthologues (Figure 8), we have not demonstrated that the changes in these cis-regulatory elements are causative. The differences in region II correlate with the up-regulation of *BnaTFL1* at the floral transition. This region is not conserved in *BnaTFL1.Cnn*, which also lacks high levels of sequence conservation in region III. The latter is associated with the expression of *AtTFL1* in Arabidopsis lateral meristems^61^ and thus predicts that *BnaTFL1.Cnn* is not expressed in this tissue.

We have shown that gene dosage and regulatory divergence may have contributed to the over-retention of flowering time genes in OSR. Without biochemical data on the proteins encoded by the genes, we are not able to distinguish whether homologues with diverged expression patterns have maintained their original molecular functions (redundant), specialised such that the initial function is split between gene duplicates (subfunctionalisation), or developed a novel function (neofunctionalisation). However, following the responsive backup circuit concept, we would expect them to have significant functional overlap.

The presence of multiple gene homologues within crop species complicates the translation of regulatory networks from models to polyploid crops, hampering breeding and selection strategies. Knowledge of functional divergence will support future breeding efforts by allowing more targeted, homologue-specific crop improvement strategies. Detailed knowledge of the function of specific copies of genes, their regulation and importantly how this functionality is combined to determine crop plasticity will be key for targeted approaches for crop improvement.

## Methods

### Plant growth and sample preparation

*Brassica napus* cv. Westar plants were sown on the 7^th^ May 2014 in cereals mix. Plants were grown in unlit glasshouses in Norwich, UK, with glasshouse temperatures set at 18 °C during the day and 15 °C at night. On the day 22, plants were transferred to a 5 °C, short day (8 hour) vernalisation room. Although Westar is classed as a spring cultivar of OSR, it may still show a mild response to the vernalisation period. After a 42-day period in the vernalisation room, plants were transferred back to unlit glasshouses and grown until the plants flowered.

The first true leaf of each plant and shoot apices were sampled at 22, 43, 64, 65, 67, 69, and 72 days after sowing (Supplementary Table 1). First true leaves were cut and immediately frozen in liquid nitrogen. The growing shoot apices were dissected using razor blades on a dry ice chilled tile before transfer to liquid nitrogen.

Samples were pooled and ground in preparation for RNA extraction. For apex tissue, ∼0.1 g of apices were ground as a pool. At the early time points, as the apices were smaller, this mass of tissue equated to approximately 20 plant apices, while at later time points approximately 10 apices were pooled. For leaf samples, between 6-10 leaf samples from separate plants were pooled and ground. RNA extraction and DNase treatment was performed following the method provided with the E.Z.N.A® Plant RNA Kit (Omega Bio-tek Inc., USA).

Library preparation and RNA sequencing was carried out by the Earlham Institute (Norwich, UK). RNA-Seq was performed on RNA samples from six time points for leaf tissue and seven time points from apex tissue. 100bp, single end reads were generated using an Illumina HiSeq2500, with an average of 67 million reads per sample (Supplementary Table 4). To assess biological variation, a second RNA sample for five time points in both the leaf and apex were sequenced at a lower average coverage of 33 million reads per sample. Supplementary Table 1 summarises the sampling scheme and indicates the time points for which a second pool of samples was sequenced.

### Gene model prediction and read alignment

Gene models are available for the Darmor-*bzh* reference genome sequence^32^ but we leveraged our sequencing data to obtain improved predictions for splice junctions. The gene model prediction software AUGUSTUS^73^ (version 3.2.2) was used to determine gene models for the Darmor-*bzh* reference genome. Tophat^74^ (version 2.0.13) aligned RNA-Seq reads from across the entire time series were combined and filtered using the `filterBam` tool provided with AUGUSTUS. AUGUSTUS used the filtered reads to aid the estimation of intron locations. Arabidopsis-derived parameters provided with the AUGUSTUS software were used to predict OSR gene models in the Darmor-*bzh* genome, with default parameters used otherwise.

RNA-Seq reads were aligned and expression levels quantified using the Tuxedo suite of software following the published workflow^75^. Tophat^74^ (version 2.0.13) with the `b2-very-sensitive`, `transcriptome-only`, and `prefilter-multihits` parameters set was used to align reads to the Darmor-*bzh* reference sequence, using the AUGUSTUS derived gene models to determine the location of gene models. Cufflinks^76^ (version 2.2.1) was used to quantify the expression levels of OSR genes. Data normalisation using `cuffnorm` was performed separately for leaf and apex tissue samples. Aside from the named parameters, default values were used.

### Identification of sequence similarity between OSR and Arabidopsis gene models

The BLAST algorithm, using the `blastn` binary provided by NCBI^77^ (version 2.2.30+) was used to identify sequence similarity between the AUGUSTUS^73^ derived gene models and the published Arabidopsis gene models downloaded from TAIR (version 10). The `blastn` algorithm was run using default parameters, with an e-value threshold of 10^-50^ used to identify sequence similarity between the AUGUSTUS derived OSR gene models and published Arabidopsis. For the analysis conducted in this study, only the most highly scoring `blastn` hit was used to identify OSR copies of Arabidopsis genes.

### Between genome expression comparison

Density plots of log_10_ transformed FPKM values were calculated and visualised using the R statistical programming language^78^. The subsets of OSR genes used showed sequence similarity to at least one published Arabidopsis gene model downloaded from TAIR^79^ (version 10), and sequence similarity to an Arabidopsis gene in the FLOR-ID database^40^ (accessed 2016-08-19).

The expression fold change for homoeologue pairs was calculated using untransformed FPKM values. The geometric mean of the fold change across all *n* homoeologous gene pairs was calculated as 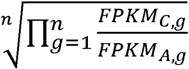 where *FPKM*_*X,g*_ is the FPKM value of the *X* genome copy of the homologue pair *g*.

### Homoeologue pair identification

The method outlined by Chalhoub et al. (2014) was used to identify pairs of homoeologues between the A and C genomes^4^. The Darmor-*bzh* reference genome was divided into the A and C genomes, removing the reference pseudo-chromosomes which consist of sequence that is unassigned to a specific chromosome. The separated genomes were uploaded to the CoGe portal^80^ and the SynMap tool^81^ was used to identify regions of syntenic genes between the two genomes. Chains of syntenic genes were identified using DAGchainer^82^, allowing a maximum 20 gene distance between two matches and with a minimum number of 4 aligned pairs constituting a syntenic block. A 1:1 synteny screen was performed using the QUOTA-ALIGN^83^ procedure. The synteny screen is necessary to distinguish homoeologous regions of the genome and paralogous regions which are the result of genome multiplication events which occurred prior to the interspecies hybridisation event in the evolutionary history of OSR. Once syntenic genes were identified using SynMap, a reciprocal sequence similarity filter was applied using the BLAST algorithm. The `blastn` algorithm was used with default parameters and a 10^-50^ e-value threshold to assess sequence similarity, and only homoeologue pairs which were reciprocal best hits in this analysis were considered. This resulted in 14427 homoeologous pairs distributed across the entire OSR genome (Supplementary Figure 9).

### Weighted gene co-expression network analysis

The weighted gene co-expression network analysis was carried out using the WGCNA library^84^ (version 1.51) available for the R statistical programming language^78^ (version 3.2.2). Due to the size of the dataset, WGCNA was performed on clustered data. The expression data was first filtered and normalised for each tissue separately. Any genes with a maximum FPKM value across the time series of less than 2.0 were removed. For the remaining genes, the expression across time was normalised to have a mean of 0.0 and a variance of 1.0. Using the normalised expression values, hierarchical clustering was conducted separately on the leaf and apex data using Euclidean distances between expression traces and a complete agglomeration method. The hierarchical tree was cut into *H* numbers of clusters and the ratio 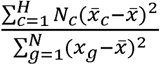 was calculated for each tree cut height, where *N* is the total number of genes, *N*_*c*_ is the total number of genes assigned to cluster *c*, *x*_*g*_ is the expression vector for gene *g*, 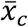 is the mean expression vector for genes assigned to cluster *c*, and 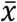 is the global mean of all expression vectors. The expression vectors are defined as 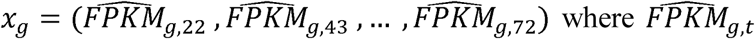 represents the normalised FPKM level of gene *g* at time point *t*, with all time points included in the vector. A ratio of ∼0.98 was chosen as a good balance between the number of clusters and how well the clusters represented the expression data. This ratio corresponded to 2683 clusters for leaf tissue and 6692 clusters for apex tissue.

WGCNA^84^ was carried out using the mean expression vectors for the 6692 apex clusters and the 2683 leaf clusters. Based on the assumption of a scale-free network structure, a soft threshold of 30 was used for both the apex and leaf samples. A minimum regulatory module size of 30 was used and modules with similar eigengene values were merged to give the final regulatory modules used for regulatory module assignment.

### Self-organising maps and the identification of regulatory modules

Self-organising maps (SOM) were generated using the `kohonen` library^85^ available for the R statistical programming language^78^. As with the WGCNA analysis, the data was filtered and normalised prior to carrying out the SOM analysis. The number of nodes used in the SOM was chosen based on the ratio 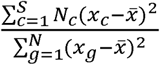 where *N* is the total number of genes, *S* is the total number of SOM nodes, *N*_*c*_ is the total number of genes assigned to SOM node *c*, *x*_*g*_ is the expression vector for gene *g*, *x*_*c*_ is the expression vector for SOM node *c*, and 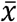 is the global mean of all expression vectors. A value of *S* was chosen such that the above ratio was ∼0.85 for both tissues. To adequately capture the variation present in the data, the dimensions of the SOM were set as the ratio between the first two principle component eigenvalues of the data, as has been done previously^86^.

To assign probabilities of genes clustering to the same SOM cluster, a resampling procedure was employed (Supplementary Figure 8a). Expression values were sampled assuming a Gaussian noise model, using the expression value as the mean of the distribution and the expression value uncertainty calculated by Cufflinks as the distribution variance. The sampled expression values for each gene, within each tissue, were normalised to a mean expression of 0.0 with a variance of 1.0 across the time series and assigned to a SOM cluster based on a minimal Euclidean distance. This sampling loop was repeated 500 times, and the SOM clusters to which the genes of interest mapped were recorded. From this process, an empirical probability of mapping to each SOM cluster was calculated for each gene of interest. The probability of two genes mapping to the same SOM cluster was then calculated as 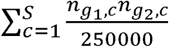 where *S* is the total number of SOM clusters, and *n*_*gi,c*_ is the number of times gene *g*_*i*_ mapped to SOM cluster *c*. As the SOM training process begins from a random starting point, some SOMs were found to better discriminate between the expression traces of some pairs of genes than other SOMs. To overcome this, the probability of two genes of interest mapping to the same SOM cluster was calculated for 100 different SOMs. This probability was averaged to give the average probability of two genes of interest mapping to the same SOM cluster.

The probability of mapping to the same cluster can also be calculated for a single gene of interest by calculating 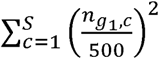. This value is a measure of how consistently a gene maps to the same SOM cluster, giving an indication of the uncertainty in the expression values calculated for that gene. Plotting a distribution of these self-clustering probabilities (Supplementary Figure 10) reveals a bimodal distribution with maxima at ∼0.05 and ∼1.0. To aid with visualising the average probabilities of two genes mapping to the same SOM cluster, as a consequence of this bimodality, a soft threshold based on a cumulative Gaussian density function was applied. The resulting value is referred to as a clustering coefficient in the main text. Clustering coefficients were calculated as 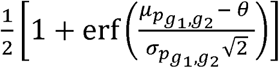 where erf is the error function defined as 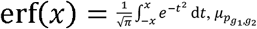 is the average probability of genes *g*_1_ and *g*_2_ mapping to the same cluster, 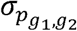 is the standard deviation of the probabilities calculated from the 100 different SOMs used in the sampling procedure, and θ is the tissue specific threshold. A threshold of 0.053 (apex) or 0.056 (leaf) was used. This threshold was calculated by taking the self-clustering probability that corresponded to the maximum of the density curve (Supplementary Figure 10) for each SOM and averaging them.

An automated approach was taken to quantify the pattern of clustering coefficients between copies of the same gene. Clustering coefficients were subjected to a binary filter, such that coefficients above 0.5 were set to 1 and those below set to 0. Regulatory modules were defined as groups of genes where the binary clustering coefficients between all genes were 1.

### Sequence conservation analysis of orthologues of *AtTFL1* in OSR

Sequence upstream and downstream of the *AtTFL1* gene was extracted from the AtGDB TAIR9/10 v171 Arabidopsis genome assembly located on PlantGDB^87^ and from the Darmor-*bzh* reference genome sequence^4^. Regions of conserved sequence were identified using mVISTA from the VISTA suite of tools^88,89^. The alignment algorithm used was AVID^90^, which performed global pair-wise alignments for all sequences. Percentage sequence conservation was calculated using a 100bp sliding window.

### Quantitative PCR of *BnaTFL1* homologues

Reverse transcription quantitative PCR (RT-qPCR) was carried out on copies of *TFL1* using custom designed primers (Supplementary Table 3). The SuperScript® III First-Strand Synthesis System (Thermo Fisher Scientific Inc., USA) was used to generate cDNA, with 2 μg of RNA used as input. The RNA was extracted as described above. Each RT-qPCR reaction consisted of 5 μl LightCycler® 480 SYBR Green I Master (Roche Molecular Systems Inc., USA), 4 μl cDNA, 0.125 μl of the forward and reverse primers at a concentration of 10 μM and 0.75 μl water. Quantification was performed on a LightCycler® 480 (Roche Molecular Systems Inc., USA). The RT-qPCR cycle consisted of a 95 °C denaturation step for 5 minutes followed by 50 quantification cycle. Each cycle consisted of 15 seconds at 95 °C, 20 seconds at 58 °C, 30 seconds at 72 °C. Fluorescence was quantified at 75 °C as the temperature was ramping from 72 °C to 95 °C.

## Data availability

All sequencing reads collected as part of this study have been made available in the NCBI Sequence Read Archive under the BioProject number PRJNA398789.

## Acknowledgements

We thank Kirsten Bomblies for critical comments on the manuscript. DMJ acknowledges support from the John Innes Foundation for the JIC Rotation PhD Programme. RW, MT, and JAI are grateful for support from BBSRC’s Institute Strategic Programme on Growth and Development (BB/J004588/1) and Genes in the Environment (BB/P013511/1). RJM acknowledges support from BBSRC’s Institute Strategic Programme on Biotic Interaction underpinning Crop Productivity (BB/J004553/1) and Plant Health (BB/P012574/1).

## Author Contributions

JAI and RJM conceived the project. DMJ, NP, RW, MT, JAI and RJM designed the experiments that were carried out by RW with support from NP, DMJ and JAI. The sequence analysis was carried out by DMJ with help from MT. DMJ performed all transcriptomic time-series analyses and produced all the figures. DMJ drafted the manuscript which was planned by DMJ, NP, JAI and RJM. All authors contributed to writing the manuscript.

